# Albumin integrates the liver’s functional status to initiate regeneration

**DOI:** 10.64898/2026.06.10.731465

**Authors:** Kristina E. Lopez, Kristin A. Knouse

**Affiliations:** Department of Biology, Massachusetts Institute of Technology, Cambridge, MA, USA; Koch Institute for Integrative Cancer Research, Massachusetts Institute of Technology, Cambridge, MA, USA

## Abstract

How an organism detects organ injury to initiate regeneration remains one of the central unresolved questions in regenerative biology. The liver is the only solid organ in mammals capable of complete regeneration, a process initiated by hepatocyte growth factor (*Hgf*) upregulation, yet the mechanism linking liver injury to *Hgf* induction has been unknown. Here we identify an albumin-based sensing mechanism that continuously reports the liver’s functional status to hepatic stellate cells to gate the regenerative program. Using single-molecule RNA FISH, parabiosis, and an *ex vivo* plasma assay, we show that stellate cells are the predominant source of *Hgf* and that a circulating, protein-dependent signal suppresses *Hgf* expression when the liver is functional. Biochemical fractionation, immunodepletion, and albumin knockout mice together demonstrate that this suppressive signal is a molecule carried by albumin rather than albumin itself. Untargeted metabolomics identified retinol as the albumin-associated suppressive molecule, which we confirm is sufficient to restore *Hgf* suppression in injured liver plasma. Conversely, long-chain fatty acids that rise after hepatectomy, specifically palmitate and linoleate, which compete for the same albumin binding site, are sufficient to derepress *Hgf* in healthy plasma. These data support a dual-input model in which albumin functions as an AND gate, integrating retinol loss and fatty acid gain as coincident indicators of liver injury before permitting *Hgf* derepression. Consistent with this model, acute *in vivo* knockdown of albumin is sufficient to derepress *Hgf* in stellate cells and drive liver overgrowth. These findings establish albumin as an active integrator of physiological signals that governs tissue-scale regenerative decisions and reveal that the liver is constitutively poised to regenerate, gatekept by signals that directly reflect its own functional status.

## INTRODUCTION

The ability to regenerate tissue after injury is observed throughout the animal kingdom, though individual species vary considerably in their regenerative capacity. For example, planaria and hydra can regenerate entire bodies, salamanders can regenerate whole limbs, and zebrafish can regenerate fins, spinal cord, heart, and liver. Decades of research have revealed insight into the early signaling events and cellular sources that drive tissue regeneration^1,2^. However, for the majority of regenerative systems, it is still unclear how the organism senses that injury has occurred and registers the need for regeneration. Understanding how organisms detect tissue damage to initiate regeneration is essential not only to achieve a complete mechanistic understanding of this process, but also for developing strategies to augment regenerative capacity in cases where it has been outstripped or in contexts where it is not currently present.

Among many mammalian species including humans and house mice, the liver stands out as the only solid organ capable of completely regenerating itself. Following liver injury, including injuries that compromise up to 75% of liver mass, the organism initiates a regenerative process that restores liver mass and function in a matter of days in rodents^3^ and weeks in humans^4^. This process is mediated by the proliferation of the liver’s differentiated cells, including the hepatocytes that constitute the bulk of liver mass and function. Hepatocyte growth factor (HGF) plays a central role in initiating this process. *Hgf* expression increases in the liver following injury^5,6^, and HGF signaling through the Met receptor is both necessary and sufficient to drive hepatocyte proliferation and liver growth^7–11^. However, as is true for other models of regeneration, it is unknown how the organism senses liver injury to upregulate *Hgf* and initiate regeneration.

## RESULTS

### Stellate cells modulate *Hgf* expression in response to a circulating signal

Although it is well-established that *Hgf* is upregulated within the liver itself upon liver injury^5,6^, the specific liver cell type(s) responsible for expressing and upregulating *Hgf* has been debated^12–15^. Recent studies increasingly nominate stellate cells as the primary cellular source of HGF^14–16^, but whether stellate cells are the only cells that upregulate *Hgf* upon liver injury remains to be conclusively demonstrated. To resolve this, we performed single-molecule RNA fluorescence *in situ* hybridization (FISH) to localize and quantify *Hgf* expression at single-cell resolution in injured and control livers. We performed two orthogonal models of liver injury: toxic injury (acetaminophen overdose) and two-thirds surgical resection (partial hepatectomy) paired with relevant controls. After 12 hours, we harvested livers and performed RNA FISH for cell type-specific markers alongside *Hgf* (Fig. 1a). In uninjured livers, *Hgf* was detected predominantly in stellate cells, with a minor contribution from sinusoidal endothelial cells and negligible expression in Kupffer cells or hepatocytes (Fig. 1b). Following both surgical resection and toxic injury, *Hgf* was exclusively upregulated in stellate cells (Fig. 1b). When considering the relative abundance of each cell type, stellate cells accounted for nearly 80% of total *Hgf* mRNA in uninjured and injured liver (Fig. 1c). Stellate cells are therefore the predominant source of *Hgf* at homeostasis and the principal cell type responsible for upregulating *Hgf* upon liver injury.

**Figure 1.**
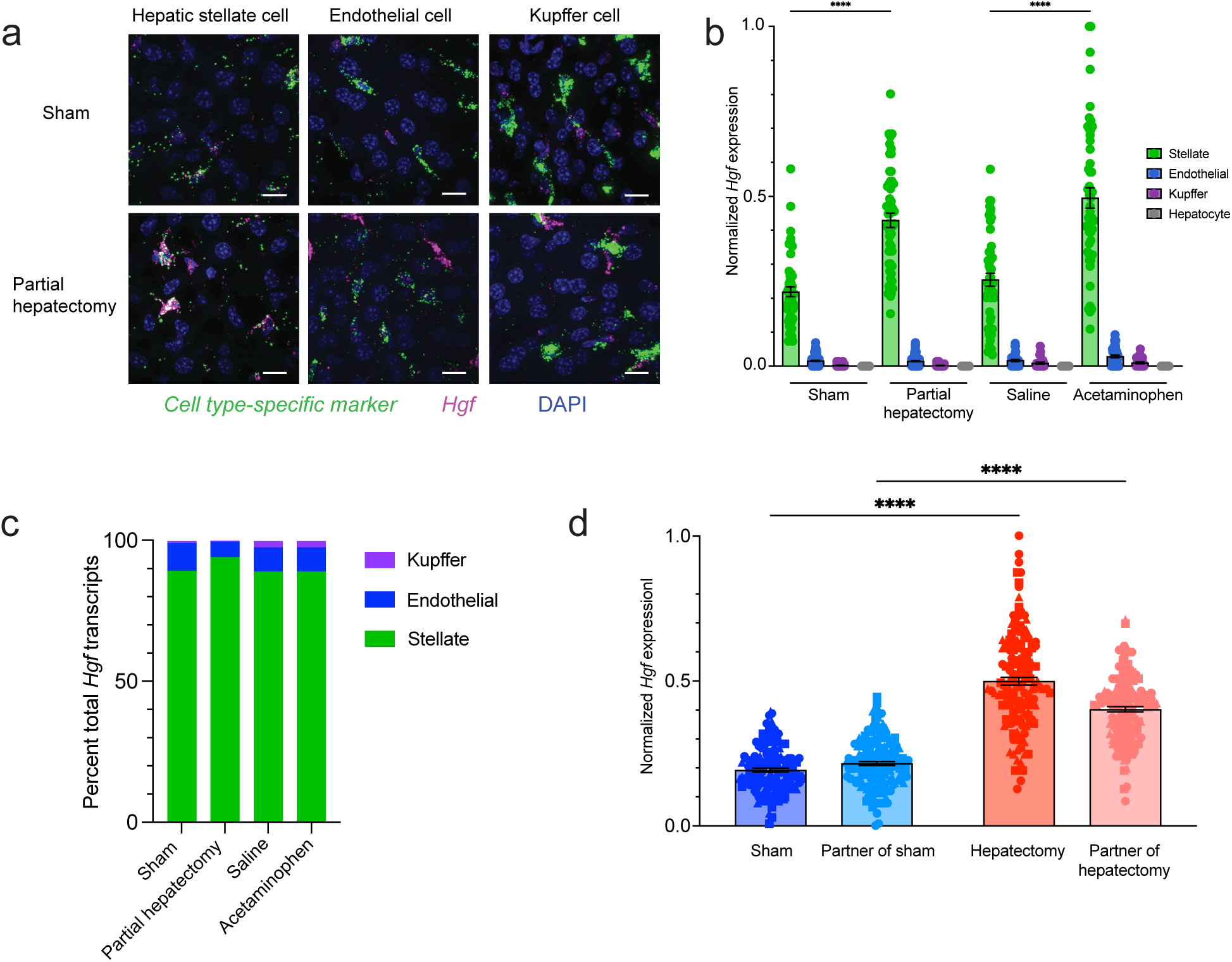
Stellate cells modulate *Hgf* in response to a circulating factor. a) RNA FISH of livers harvested 12 hours after sham surgery or partial hepatectomy stained for a cell type-specific mRNA (*Lrat* for stellate cells, *Aqp1* for endothelial cells, or *Clec4f* for Kupffer cells) and *Hgf*, and counterstained with DAPI. Scale bar, 10 μm. b) Normalized *Hgf* expression in each cell type 12 hours after sham surgery, partial hepatectomy, saline, or acetaminophen as determined by RNA FISH. n = 3 male and 3 female mice per condition with ≥ 25 cells of each type per mouse. Data are mean ± s.e.m. Sham versus partial hepatectomy stellate cells p <0.0001, saline versus acetaminophen stellate cells p < 0.0001 by one-way ANOVA with Tukey’s multiple comparisons test. c) Percent of total *Hgf* transcripts in the liver contributed by each cell type. d) Normalized *Hgf* expression per stellate cell 24 hours after partial hepatectomy or sham surgery in parabiotic pairs, as determined by RNA FISH. n = 3 mouse pairs per perturbation. 50 cells quantified per mouse. Data are mean ± s.e.m. Sham versus hepatectomy p < 0.0001, partner of sham versus partner of hepatectomy p < 0.0001 by one-way ANOVA with Tukey’s multiple comparisons test.

Having established that stellate cells are the cells responsible for upregulating *Hgf* after injury, we began to investigate how stellate cells detect that liver injury has occurred. We first sought to determine whether *Hgf* regulation is mediated by a circulating factor in the bloodstream or by a local signal within the liver. To answer this, we performed parabiosis on pairs of mice and then performed either sham surgeries or partial hepatectomies on one mouse of each pair while leaving its partner unperturbed. We confirmed that the parabiotic pairs had established shared circulation by observing elevated serum alanine aminotransferase (ALT) and aspartate aminotransferase (AST), enzymes released by the hepatectomized liver^17^, in the partners of hepatectomized mice (Extended Data Fig. 1a,b). We then investigated *Hgf* expression in the stellate cells of both perturbed animals and their partners using RNA FISH. Notably, in the setting of partial hepatectomy, stellate cells in the uninjured partner of the hepatectomized mouse also upregulated *Hgf* (Fig. 1d). We therefore conclude that stellate cells modulate *Hgf* expression in response to a circulating signal.

### An albumin-dependent signal suppresses *Hgf* at homeostasis

To facilitate further characterization of the circulating signal that regulates *Hgf* expression, we asked whether we could recapitulate this signaling *ex vivo* using mouse plasma and primary stellate cells. We isolated non-parenchymal cells (including stellate cells, endothelial cells, and Kupffer cells) from unperturbed mice, cultured them for three hours in plasma harvested from separate mice, and quantified *Hgf* expression by RNA FISH. Stellate cells cultured in plasma from mice that had undergone a partial hepatectomy (hepatectomy plasma) had increased *Hgf* expression relative to cells cultured in either plasma from mice receiving a sham surgery (sham plasma) or plasma from unperturbed mice (Fig. 2a). This established that we could indeed recapitulate this signaling *ex vivo* and confirmed that a factor in mouse plasma regulates *Hgf* expression.

**Figure 2.**
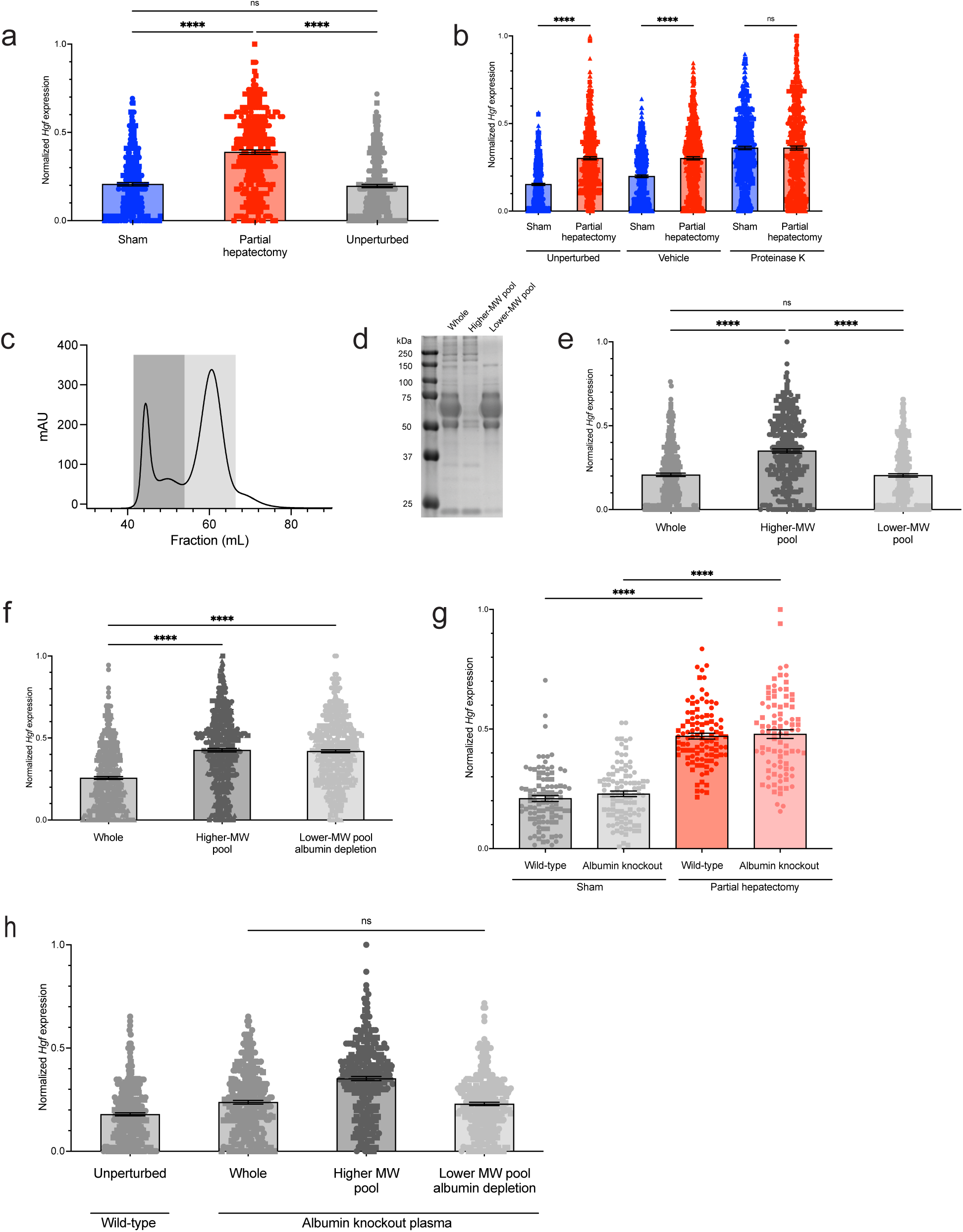
An albumin-dependent signal suppresses *Hgf* at homeostasis. a) Normalized *Hgf* expression in stellate cells as determined by RNA FISH. n = 3 biological replicates with 150 cells analyzed per condition per replicate. Data are mean ± s.e.m. Unperturbed versus hepatectomy p < 0.0001, sham versus hepatectomy p < 0.0001 by one-way ANOVA with Tukey’s multiple comparisons test. b) Normalized *Hgf* expression in stellate cells as determined by RNA FISH. n = 3 biological replicates with 150 cells analyzed per condition per replicate as determined by RNA FISH. Data are mean ± s.e.m. Untreated sham versus untreated hepatectomy p < 0.0001, vehicle sham versus vehicle hepatectomy p < 0.0001 by one-way ANOVA with Tukey’s multiple comparisons test. c) Absorbance measurements across fractions upon size-exclusion chromatography of unperturbed plasma. Dark grey box represents the higher-molecular weight pool, and light grey box represents lower-molecular weight pool. d) Coomassie-stained SDS-PAGE gel of equal input of whole plasma, higher-, and lower-molecular weight (MW) pools. e) Normalized *Hgf* expression in stellate cells as determined by RNA FISH. n = 2 biological replicates with 150 cells analyzed per condition per replicate. Data are mean ± s.e.m. Higher-MW versus lower-MW pool p < 0.0001 by one-way ANOVA with Tukey’s multiple comparisons test. f) Normalized *Hgf* expression in stellate cells as determined by RNA FISH. n = 3 biological replicates with 150 cells analyzed per condition per replicate. Data are mean ± s.e.m. Whole versus higher-MW pool p < 0.0001 and whole versus lower-MW pool, albumin depleted p < 0.0001 by one-way ANOVA with Tukey’s multiple comparisons test. g) Normalized *Hgf* expression in stellate cells as determined by RNA FISH. n = 2 biological replicates with 50 cells analyzed per condition per replicate. Data are mean ± s.e.m. Wild-type sham versus hepatectomy p < 0.0001, albumin knockout sham versus hepatectomy p < 0.0001 by one-way ANOVA with Tukey’s multiple comparisons test. h) Normalized *Hgf* expression in stellate cells as determined by RNA FISH. n = 2 biological replicates with 150 cells analyzed per condition per replicate. Data are mean ± s.e.m.

To determine whether the factor regulating *Hgf* expression is protein-dependent, we treated sham and hepatectomy plasma with thermolabile proteinase K to degrade proteins prior to culturing with stellate cells. Proteinase K treatment of sham plasma caused stellate cells to upregulate *Hgf* to a level equivalent to that of hepatectomy plasma (Fig. 2b). This result suggests that the default state of stellate cells is to express high levels of *Hgf*, and that a protein-dependent factor suppresses *Hgf* expression when the liver is functional.

To narrow the identity of the protein-dependent suppressive signal, we performed size-exclusion chromatography on plasma from unperturbed mice, resolving it into higher- and lower-molecular weight pools, and treated stellate cells with these two pools (Fig. 2c,d). Suppressive activity was retained exclusively in the lower-molecular weight pool of plasma from unperturbed mice (Fig. 2e). Similar fractionation experiments using hepatectomy plasma showed that neither pool had suppressive activity (Extended Data Fig. 2a). These data reaffirm that a protein-dependent signal suppresses *Hgf* expression when the liver is functional and narrowed the signal to the lower-molecular weight pool.

Albumin, the most abundant plasma protein, was among the proteins in the lower-molecular weight pool (Fig. 2d). Given its abundance, we decided to test whether albumin is required to suppress *Hgf* expression. We immunodepleted albumin from the lower-molecular weight pool of unperturbed plasma (Extended Data Fig. 2b). Albumin depletion abolished the suppressive activity of the lower-molecular weight pool, indicating that albumin is required to suppress *Hgf* expression when the liver is functional (Fig. 2f).

Albumin serves two main functions in plasma. First, by virtue of its high concentration, it maintains plasma oncotic pressure. Second, by virtue of its ability to bind various molecules, it carries molecules that would otherwise be insoluble in aqueous plasma. We therefore hypothesized that albumin-dependent suppression of *Hgf* expression in stellate cells could occur via albumin itself, a post-translational modification of albumin, or a molecule carried by albumin. We noted that plasma concentrations of albumin were not significantly different twelve hours after partial hepatectomy relative to sham (Extended Data Fig. 2c), arguing against concentration-dependent signaling from albumin itself and instead focusing our attention on a post-translational modification of albumin or a molecule carried by albumin.

To distinguish between a post-translational modification and a carried molecule as the albumin-dependent suppressive signal, we turned to human and mouse genetics. Both humans and mice with homozygous loss-of-function mutations in albumin exist and are largely healthy^18,19^. Studies of these individuals’ plasma shows upregulation of other plasma proteins, which is believed to compensate for the oncotic and carrier functions of albumin^18,19^. If the suppressive signal were albumin itself or a post-translational modification of albumin, albumin knockout mice should be unable to regulate *Hgf* appropriately. Conversely, if the signal were a molecule carried by albumin, it may transfer to one of the compensatory carrier proteins. We obtained albumin knockout mice and confirmed absence of albumin in their plasma (Extended Data Fig. 3a). Albumin knockout mice had normal liver size at baseline (Extended Data Fig. 3b). Further, albumin knockout mice had equivalent *Hgf* expression to wild-type mice in the absence of liver injury and appropriately upregulated *Hgf* after partial hepatectomy (Fig. 2g). These data argue against the suppressive signal being albumin itself or a post-translational modification of albumin and instead suggest that the signal is a molecule normally carried by albumin that transfers to a compensatory carrier protein in its absence. To further test this, we asked whether albumin knockout plasma could still suppress *Hgf* expression in wild-type stellate cells. Indeed, the lower-molecular weight fraction of albumin knockout plasma retained suppressive activity in wild-type stellate cells even after mock albumin depletion (Fig. 2h). Together, these data suggest that a molecule normally carried by albumin suppresses *Hgf* in stellate cells and are consistent with a model in which liver injury provokes loss of this signal which in turn leads to derepression of *Hgf*.

### Retinol and long-chain fatty acids antagonistically regulate *Hgf* expression

We next sought to identify the albumin-dependent signal and understand how this signal is lost upon liver injury to derepress *Hgf.* We reasoned that the albumin-associated levels of this signal should be relatively high at homeostasis and reduced upon liver injury. Further, this signal should be present in uninjured albumin knockout plasma but absent in uninjured wild-type plasma depleted of albumin. In principle, liver injury could either reduce the overall levels of this molecule and/or directly displace this molecule from albumin. To identify candidate suppressive signals and potential displacing molecules, we performed untargeted metabolomics on multiple replicates of plasma from unperturbed, sham, or hepatectomized wild-type mice, their corresponding lower-molecular weight fractions, the albumin-depleted lower-molecular weight fraction of wild-type plasma, and the lower-molecular weight fraction of albumin knockout plasma (Extended Data Fig. 4a). This identified 929 metabolites across 72 samples. We confirmed expected changes in metabolites known to be affected by liver injury, including increased bilirubin and decreased glucose in the hepatectomy samples as compared to sham and unperturbed samples (Extended Data Fig. 4b,c). Additionally, two molecules established to be carried by albumin, tryptophan and thyroxine, were present in the lower-molecular weight pools of unperturbed, sham, and hepatectomy plasma and depleted upon albumin immunodepletion (Extended Data Figs. 4d,e), confirming our ability to detect albumin-associated metabolites.

We applied two complementary criteria to identify candidate suppressive signals: enrichment in the lower-molecular weight pools of sham relative to hepatectomy plasma, and depletion following albumin immunodepletion in wild-type plasma relative to albumin knockout plasma. Seven metabolites satisfied both criteria: glutamate, aspartate, retinol, leucine, tryptophan, N-formylanthranilic acid, and arginine (Fig. 3a,b). Of these, only retinol, tryptophan, and N-formylanthranilic acid are believed to require albumin for transport in plasma^20^. Of these, retinol and tryptophan were particularly intriguing because vitamin A metabolism and tryptophan homeostasis are tightly coupled to hepatocyte function, making them plausible indicators of liver functional status.

**Figure 3.**
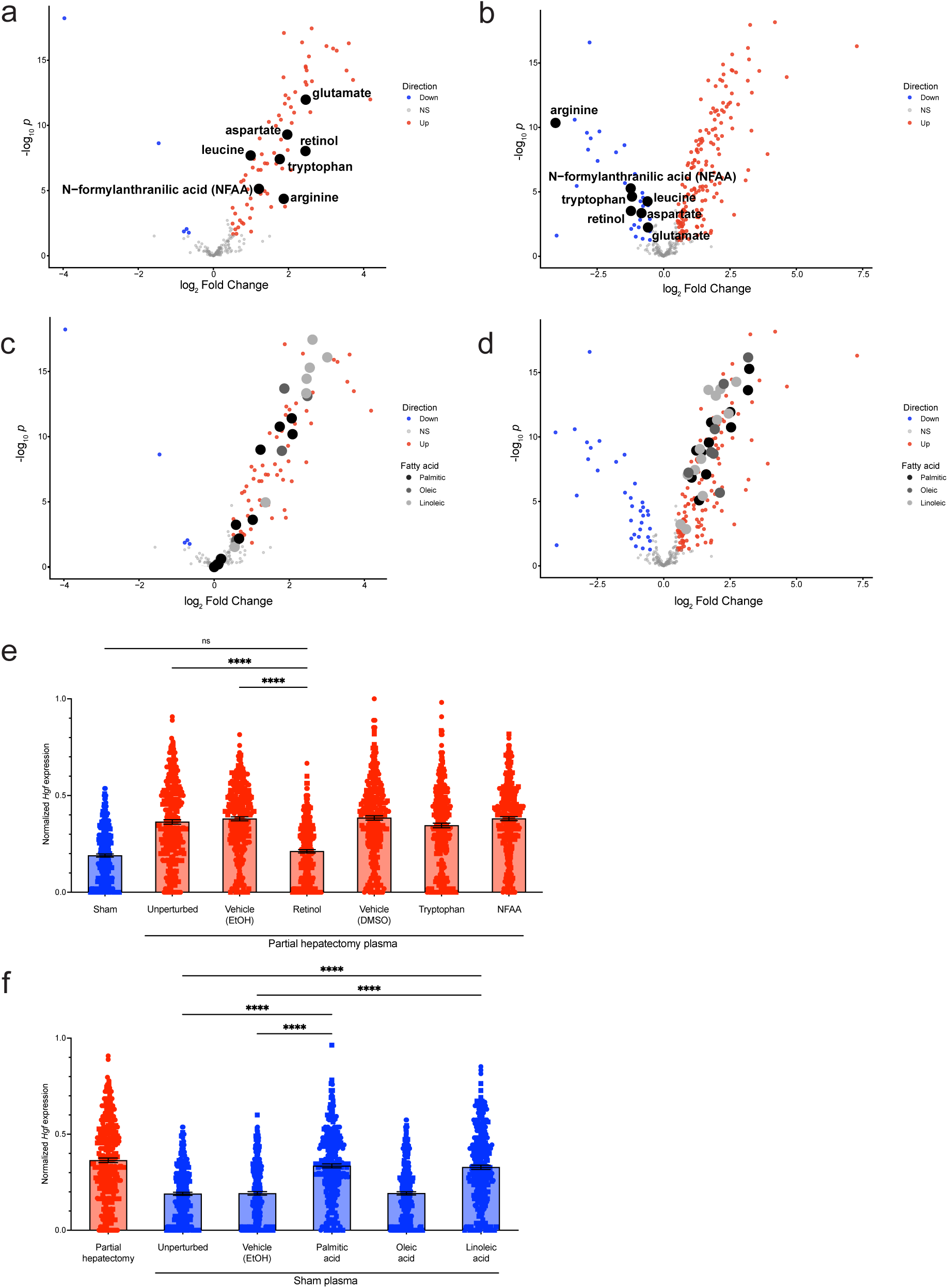
Retinol and long-chain fatty acids antagonistically regulate *Hgf* expression. a) Differentially abundant metabolites in the lower-molecular weight fraction of hepatectomy plasma compared to lower-molecular weight fraction of sham plasma. Metabolites highlighted in black indicate high-priority candidates. b) Differentially abundant metabolites in the lower-molecular weight fraction of albumin knockout plasma compared to lower-molecular weight fraction of wild-type plasma after albumin depletion. Metabolites highlighted in black indicate high-priority candidates. c) Differentially abundant metabolites in the lower-molecular weight fraction of hepatectomy plasma compared to lower-molecular weight fraction of sham plasma. Metabolites highlighted indicate potential displacing molecules. d) Differentially abundant metabolites in the lower-molecular weight fraction of albumin knockout plasma compared to lower-molecular weight fraction of wild-type plasma after albumin depletion. Metabolites highlighted indicate potential displacing molecules. e) Normalized *Hgf* expression in stellate cells as determined by RNA FISH. n = 2 biological replicates with 150 cells analyzed per condition per replicate. Data are mean ± s.e.m. Hepatectomy plasma with retinol vehicle (EtOH) versus hepatectomy plasma with retinol p < 0.0001 by one-way ANOVA with Tukey’s multiple comparisons test. f) Normalized *Hgf* expression in stellate cells as determined by RNA FISH. n = 2 biological replicates with 150 cells analyzed per condition per replicate. Data are mean ± s.e.m. Sham plasma with fatty acid vehicle (EtOH) versus sham plasma with palmitic acid p < 0.0001, sham plasma with fatty acid vehicle (EtOH) versus sham plasma with linoleic acid p < 0.0001 by one-way ANOVA with Tukey’s multiple comparisons test.

We also considered the possibility that another molecule(s) displaces the suppressive signal from albumin upon liver injury. To identify candidate displacing molecules, we identified molecules that were enriched in the hepatectomy relative to sham lower-molecular weight pool and depleted following albumin immunodepletion of wild-type plasma relative to albumin knockout plasma. Several long-chain fatty acids, including palmitic acid, oleic acid, linoleic acid, and derivatives of these fatty acids, fit the criteria as candidate displacers (Fig. 3c,d). Long-chain fatty acids were intriguing candidate displacers since circulating fatty acids are known to increase following partial hepatectomy.

To determine whether any of the candidate signals were sufficient to suppress *Hgf*, we supplemented hepatectomy plasma with supraphysiologic concentrations of retinol, tryptophan, and N-formylanthranilic acid and then evaluated its effects on *Hgf* expression in stellate cells. Interestingly, supplementing hepatectomy plasma with retinol, but none of the other molecules, was sufficient to suppress *Hgf* to levels equivalent to sham plasma (Fig. 3e). These data implicate retinol as the albumin-dependent signal that suppresses *Hgf* when the liver is functional.

Retinol is a lipophilic molecule established to bind albumin at fatty acid site 1^21^, which is also a primary binding site for many long-chain fatty acids. This increased our suspicion that long-chain fatty acids might displace retinol from albumin upon liver injury. To test whether long-chain fatty acids can derepress *Hgf*, we supplemented sham plasma with supraphysiologic concentrations of three different long-chain fatty acids: palmitic acid, known to bind fatty acid sites 1, 2, and 7; oleic acid, known to bind fatty acid sites 3, 4, and 5; and linoleic acid, known to bind fatty acid sites 2, 4, and 5. Notably, palmitic acid and linoleic acid, but not oleic acid, were sufficient to derepress *Hgf* to levels equivalent to hepatectomy plasma (Fig. 3f). The ability of linoleic acid to suppress *Hgf* expression despite not binding to fatty acid site 1 could be explained by the well-established cooperative nature of albumin binding, in which ligand binding at one site can influence ligand affinity at others^22^. Collectively, these data support a model in which retinol, present at a high concentration in plasma at homeostasis, serves to suppress *Hgf* expression in stellate cells. Upon liver injury, a drop in plasma levels of retinol coupled to an increase in plasma levels of palmitic and linoleic acid cooperate to trigger a dramatic reduction in the levels of albumin-associated retinol, leading to derepression of *Hgf*.

### Acute albumin knockout derepresses *Hgf in vivo*

Our *ex vivo* data indicate that albumin is essential for mediating *Hgf* suppression. Although we find that albumin is dispensable in the setting of constitutive albumin knockout, presumably due to the upregulation of compensatory carrier proteins, we wondered whether acute depletion of albumin would not be effectively compensated and thereby lead to derepression of *Hgf in vivo*. To test this, we acutely knocked out albumin in adult mice using an inducible CRISPR system. We generated liver-targeting adeno-associated viruses (AAV) expressing Cre and sgRNA targeting either *Albumin* or the non-coding *Tigre* locus and injected these AAV vectors into Cre-inducible Cas9 adult mice to achieve liver-wide knockout. Although we never achieved complete depletion of albumin using this system, we found that by 56 days after injection plasma levels of albumin were substantially reduced in the sgAlbumin mice relative to sgTigre mice. (Fig. 4a,b). Notably, acute albumin knockout led to a derepression of *Hgf* in both male and female mice (Fig. 4c). This derepression was accompanied by a significant increase in liver-to-body weight ratio in males but not females at 56 days after AAV injection (Fig. 4d). No significant changes in other organ weights were observed at this timepoint (Extended Data Fig. 5a-e). The liver overgrowth phenotype in males became more pronounced by 112 days, with liver mass nearly doubling relative to *Tigre* controls (Fig. 4e,f). Female albumin knockout mice again showed no significant liver enlargement (Fig. 4e,f), revealing an unexpected sex-specific difference in the downstream growth response to *Hgf* derepression. Collectively, these data support our *ex vivo* findings that albumin is required to suppress *Hgf* expression at homeostasis. Further, we find that acute depletion of albumin is sufficient to derepress *Hgf in vivo* and, in male mice, trigger liver overgrowth.

**Figure 4.**
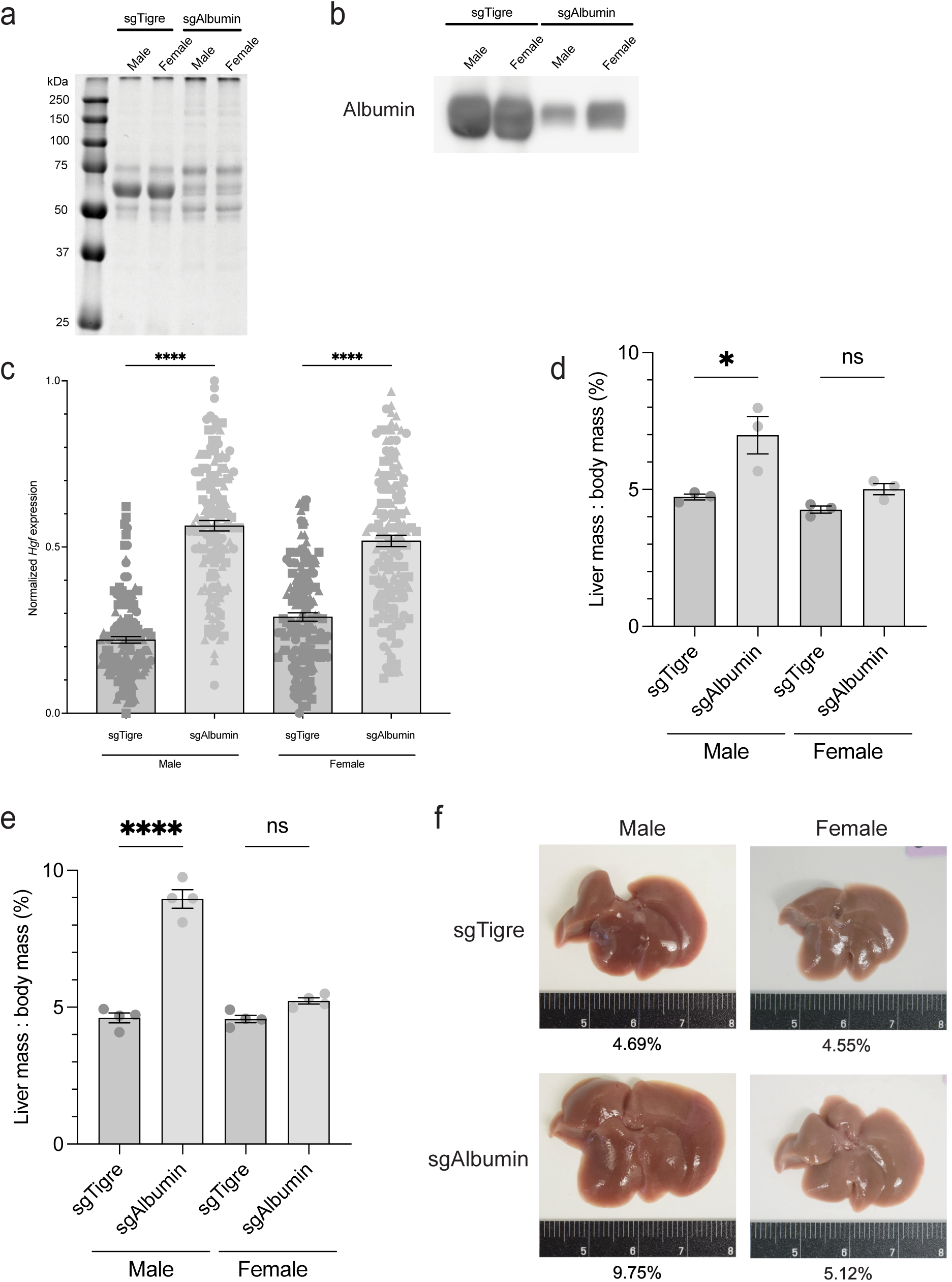
Acute albumin knockout derepresses *Hgf in vivo*. a) Coomassie-stained SDS-PAGE gel of equal input of plasma harvested 56 days after injection with AAV-Cre-sgTigre or AAV-Cre-sgAlbumin b) Western blot of equal input of plasma harvested 56 days after injection with AAV-Cre-sgTigre or AAV-Cre-sgAlbumin. c) Normalized *Hgf* expression in stellate cells as determined by RNA FISH. n = 3 biological replicates with 50 cells analyzed per condition per replicate. Data are mean ± s.e.m. sgTigre male versus sgAlbumin male p < 0.0001, sgTigre female versus sgAlbumin female p < 0.0001 by one-way ANOVA with Tukey’s multiple comparisons test. d) Liver-to-body weight ratio 56 days after injection with AAV-Cre-sgTigre or AAV-Cre-sgAlbumin. Data are mean ± s.e.m. e) Liver-to-body weight ratio 112 days after injection with AAV-Cre-sgTigre or AAV-Cre-sgAlbumin. Data are mean ± s.e.m. f) Images of livers 112 days after injection with AAV-Cre-sgTigre or AAV-Cre-sgAlbumin. Ruler for scale. Liver-to-body weight ratio (%) listed below each picture.

## DISCUSSION

Our findings bridge a mechanistic gap in our understanding of liver regeneration by identifying stellate cells as the source of injury-induced *Hgf* and establishing a mechanism by which liver function is communicated to stellate cells to regulate *Hgf* expression. We specifically find that albumin-associated retinol suppresses *Hgf* in stellate cells at homeostasis. Following liver injury, a drop in the plasma concentration of retinol coupled to an increase in long-chain fatty acids may collectively act to reduce albumin-associated retinol availability and thereby relieve *Hgf* suppression. We show that acute loss of albumin *in vivo* is sufficient to derepress *Hgf* in stellate cells and drive liver overgrowth, establishing albumin as an essential mediator in the communication of liver function and initiation of regeneration.

In principle, initiation of a regenerative program could be mediated either by the appearance of an activating signal or by the loss of a suppressive signal. Further, the precise identities of these signals could originate from the injury itself or the ensuing loss of organ function. For example, in skin, there is evidence that wound-induced hypoxia directly triggers upregulation of pro-regeneration cytokines, a model in which the injury itself actively triggers regeneration^23^. Our findings reveal a fundamentally different logic in the liver. We find that the default state of stellate cells is to express high levels of the pro-regenerative factor *Hgf*, and this is suppressed at baseline by the liver-derived metabolite retinol. The liver is thus poised to regenerate at baseline and held in check by signals that directly reflect liver function.

Our data support a model in which albumin integrates the liver’s functional status and communicates this status to stellate cells to determine whether or not regeneration should occur. When the liver is functional, albumin-associated retinol levels are high and stellate cells suppress *Hgf*. When the liver is injured, long-chain fatty acids rise in the plasma. These long-chain fatty acids can displace retinol from albumin to derepress *Hgf*. According to this model, albumin functions as a physiological “AND” gate, integrating two orthogonal indicators of liver function before permitting activation of the regenerative program. This dual-input logic can confer robustness to the system, ensuring that the regenerative program is not inappropriately engaged by fluctuations in either signal alone. Our data thus highlight albumin as a protein capable of integrating multiple signals and communicating these signals to responding cells, thereby enabling robust regulation of physiology. How exactly albumin-associated retinol signals to stellate cells is an important future direction.

Albumin’s central role in integrating signals of liver function and communicating these signals to stellate cells to control regeneration reveals albumin to be a novel therapeutic target for modulating liver regeneration. A major unmet need in clinical hepatology is a treatment for acute liver failure, a clinical scenario in which extreme liver injury outstrips the liver’s regenerative capacity. Notably, it’s been shown that ectopic administration of HGF can improve recovery and survival in acute liver failure^24^, however the extremely short half-life of HGF hinders the feasibility of such therapy^25^. Our data raise the exciting possibility of pharmacologically displacing retinol from albumin to augment *Hgf* expression from its natural source. More broadly, our data highlight that albumin is not merely a passive carrier protein but rather an active integrator of physiological signals that govern tissue-scale decisions, a function that may extend to other albumin-bound molecules and other tissues in which albumin-dependent signaling has yet to be appreciated.

## METHODS

### Partial hepatectomy

Two-thirds partial hepatectomy was performed on 8–12-week-old mice as previously described^26^. Briefly, 8–12-week-old male mice were anesthetized with 2% isoflurane-oxygen delivered via nosecone. Pre-operative analgesia in the form of subcutaneous carprofen (Rimadyl) at a dose of 5 mg/kg was administered subcutaneously. Fur on the abdomen was shaved, and the region was sterilized with betadine and 70% isopropanol in succession three times. A midline incision was made from the midpoint of the abdomen to the sternum. An Alm retractor was placed beneath skin and peritoneum and expanded to reveal liver. Different 4-0 silk sutures were tied at the base of the right median lobe, left median lobe, and left lateral lobes sequentially. After tying of each knot, the lobe was cut and removed. The abdomen was washed with 1 mL of warmed lactated ringers’ solution and excess was absorbed with sterile gauze. The peritoneum was sutured closed using an absorbable 5-0 vicryl suture. The overlying skin was closed using 7 mm wound clips. The mouse was then given 20 mL/kg of warmed lactated ringers’ solution subcutaneously and placed on a heating pad to recover. 90% partial hepatectomies were performed as described in the 70% partial hepatectomy description above, except the right inferior lateral lobe was also removed. Mice were sacrificed at 12 or 24 hours following surgery.

### Acetaminophen injury

Acetaminophen was dissolved in warm 0.9% saline and injected intraperitoneally at 300 mg/kg into 8–12-week-old mice after a 12 hour fast. Mice were sacrificed at 12 or 24 hours following injection.

### Parabiosis

Parabiosis surgeries were performed as previously described^27^. Briefly, 6-week-old female mice were housed in weight matched pairs (within 1 g) for two days prior to surgery. Both mice were anesthetized with 2% isoflurane-oxygen in the induction chamber, once down, the mice were moved to a double outlet nosecone. Pre-operative analgesia was administered Bup-SR at 1 mg/kg and carprofen (Rimadyl) at 5 mg/kg, both subcutaneously. Each mouse was shaved from under their ear to their hind leg, going from near their spine down toward the midline of their abdomen. Nair was used to remove any remaining hair from the shaved region, and 70% isopropanol was used to remove all Nair. The area was sterilized with alternating betadine and 70% isopropanol covered gauze three times. A longitudinal incision was made in the skin from the thigh to just below the ear of each mouse. Then, an incision in the peritoneum was made from above the adipose tissue above the thigh to the rib cage. The peritoneums of mice were then joined using a continuous absorbable 5-0 PDS-II suture in a closed fashion, such that there is no hole for the intestines to cross into the other mouse. A 3-0 silk suture was used to join the muscle over the shoulder blades of each mouse and tied with a knot. To close the skin, a continuous suture was done using a 5-0 PDS-II suture, starting two-thirds down the mouse’s back, and done until the same spot was reached, tying off with a double knot. Mice received 20 mL/kg of warmed lactated ringers’ solution subcutaneously and were placed on a new cage on heat. The first three days following surgery, each mouse was given 5 mg/kg Carprofen and 20 mL/kg of warmed lactated ringers.

### Non-parenchymal cell isolation

Non-parenchymal cells were isolated from mice using a pronase-collagenase IV digestion, based on a previously published protocol^28^, with modifications. 8–12-week-old male mice were anesthetized with 2% isoflurane-oxygen delivered via nosecone. The liver was perfused with 30 mL of warm EGTA solution by incising the portal vein with a 25G needle attached to a peristaltic pump. The inferior vena cava was incised immediately after perfusion began to allow fluid outflow. Following EDTA perfusion, the liver was cut into thin slices and placed into a flask with the pronase and collagenase IV digestion solution. The cell-enzyme solution was placed into a 37°C shaking water bath to shake at 174 RPM for 30 minutes. Following the *ex vivo* digestion, the cell solution was filtered through a 40 μm cell strainer into a 50 mL conical and spun at 50 G for 5 minutes at 4°C to pellet any hepatocytes. Then, the supernatant was transferred to a clean 50 mL and spun at 580 G for 10 minutes at 4°C to pellet all non-parenchymal cells. Supernatant was aspirated down to 10 mL then 100 μL DNase was added and the pellet was gently resuspended. The conical was then filled to 50 mL with GBSS and spun again at 580 G for 10 minutes at 4°C. Supernatant was then fully aspirated, 1mL of GBSS and 100 μL DNase was used to resuspend the final pellet and then cells were used in downstream assays.

### Single-molecule RNA fluorescence *in situ* hybridization of tissues

Liver cross-sections of approximately 3 mm were fixed in 4% paraformaldehyde at 4°C for three hours, then fixed in a 4% paraformaldehyde and 30% sucrose solution overnight. Livers were then embedded in OCT and sectioned to 10 μm. Slides were airdried at -20°C for 1 hour. Slides were washed in PBS for 5 minutes to remove OCT, followed by incubation at 60°C for 30 minutes. Slides were then immersed in 4% paraformaldehyde for 1 hour at room temperature. Slides were dehydrated by immersion in 50%, 70%, 100%, 100% EtOH, each for 5 minutes, then slides were airdried. RNAscope Hydrogen Peroxide solution was dropped over the tissue and incubated for 10 minutes before two water washes to remove the hydrogen peroxide. Slides were then boiled in 1x RNAscope Target Retrieval solution at 97°C for 5 minutes. Immediately after boiling, slides were submerged in excess water and washed three times. Then, slides were again washed in 100% EtOH before airdrying overnight. After pretreatment, the manufacturer’s protocol for the RNA Multiplex Fluorescent Detection Kit v2 was followed. Advanced Cell Diagnostics designed all probes used in this study (*Hgf, Lrat, Aqp1, Clec4f*). RNAscope was performed on livers from 3 mice per condition. Images were acquired using the TissueFAXS SL Fluorescent Slide Scanner using a 40X air objective. Images were analyzed using the TissueFAXS Viewer software. 50 stellate cells per liver per condition were analyzed. For visualization and comparison across experiments, values were min-max normalized according to:

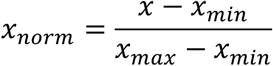

where *x*_min_ and *x*_max_ represent the minimum and maximum values observed within the indicated dataset, respectively. Each dataset included one complete cohort of conditions.

### Single-molecule RNA fluorescence *in situ* hybridization of single cells

Single cells were fixed in 4% paraformaldehyde for 30 minutes with rocking at room temperature. Cells were spun down at 580 G at 4°C for 10 minutes, washed with PBS, and again spun down 580 G at 4°C for 10 minutes. PBS was aspirated and cells were resuspended in 70 μL of PBS and dropped onto slides and left to dry overnight at room temperature. The following day, the slides were dehydrated by immersion in 50%, 70%, 100%, 100% EtOH, each for 5 minutes. The slides were then stored in 100% EtOH at - 20°C overnight or up to one month. When starting the RNAscope protocol, the slides were dried at 37°C for 30 minutes. After this, the manufacturer’s protocol for the RNA Multiplex Fluorescent Detection Kit v2 was followed. Advanced Cell Diagnostics designed all probes used in this study (*Hgf, Lrat, Aqp1, Clec4f*). RNAscope was performed on fixed cells from 3 mice per condition. Images were acquired using the TissueFAXS SL Fluorescent Slide Scanner using a 40X air objective. Images were analyzed using the TissueFAXS Viewer software. 150 stellate cells per replicate per condition were analyzed. For visualization and comparison across experiments, values were min-max normalized according to:

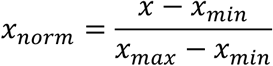

where *x*_min_ and *x*_max_ represent the minimum and maximum values observed within the indicated dataset, respectively. Each dataset included one complete cohort of conditions.

### Blood harvest

Mice were anesthetized with 2% isoflurane-oxygen delivered via nosecone. A “U-shaped” incision was made from the base of the abdomen to the ribcage. The portal vein was incised using a 25G needle attached to a 1 mL syringe, with the needle pointed inferior to the liver and blood was collected. For plasma, blood was placed in EDTA K3E coated tubes from Sarstedt (Cat.# 41.1395.105) and spun at 3500 G for 6 minutes at room temperature. For serum, blood was placed in BD Microtainer® Blood Collection Tubes (Cat.# 365967) and allowed to clot for 30 minutes at room temperature, followed by a 10400 G spin for 2 minutes at room temperature.

### Serum ALT and AST measurements

Serum isolated from mice as described above. Samples were submitted to IDEXX for measurements of ALT and AST.

### Proteinase K treatment of plasma

Plasma was harvested from sham-operated or partially hepatectomized mice. Aliquots of 250 μL was incubated with 50 μL of vehicle control (20 mM Tris-HCl, 1 mM CaCl2, 50% Glycerol, pH 7.4 @ 25°C) or 50 μL of Proteinase K (Cat.# P8111S) and left at room temperature for 6 hours. To inactivate the Proteinase K, the plasma was incubated at 55°C for 10 minutes then used in downstream assays.

### Size-exclusion chromatography

Size-exclusion chromatography was performed using the AKTA Pure FPLC system and a HiPrep 16/60 Sephacryl S-200 HR Column. The column was washed with water at a rate of 0.5 mL/min for three column volumes (360 mL). The column was then equilibrated with PBS at a rate of 0.5 mL/min for three column volumes (360 mL). Once ready, 500 μL of filtered plasma was injected into the AKTA system and ran on the column at a rate of 1 mL/min, with a fraction volume set to 1 mL. After 94 mL, fractions were collected. Fractions pertaining to the higher-molecular weight peak (39-50) and lower-molecular weight peak (51-62) were concentrated using Amicon® Ultra Centrifugal Filters with a 10 kDa MWCO (Cat.# UFC8010). After concentration, volumes for each pool were brought up to 250 μL, then used for downstream assays.

### Immunodepletion of albumin

Albumin was immunodepleted from the lower-molecular weight fraction pool using the Proteome Purify 2 Mouse Serum Protein Immunodepletion Resin (Cat.# MIDR002) per manufacturer’s instructions. Briefly, following concentration, the lower-molecular weight pool was brought up to 400 μL with PBS. 10 μL of plasma was aliquoted in separate 12 mL tubes and 1 mL of resin was added to each tube. The tubes were placed on a rotating spinner for one hour. To remove the resin, the liquid in each tube was placed in a Spin-X Filter Unit (Cat.# 8160) and spun down at 2000 G for 2 minutes. The flowthrough was collected and concentrated using an Amicon® Ultra Centrifugal Filters with a 10 kDa MWCO. Volume was brought up to 250 μL, then used for downstream assays.

### Albumin ELISA

Albumin ELISAs were performed using an Albumin ELISA Kit (Cat.# OKIA00087) per manufacturer’s instructions. Briefly, plasma was diluted 1 to 500,000 then aliquoted into separate wells of the pre-coated plate to form Albumin + Anti-Albumin complexes. Four washes were performed to remove unbound plasma proteins. The Anti-Albumin-HRP conjugate was added, then four more washes performed to remove unbound conjugate. Finally, the chromogenic substrate was added and the absorbance at 450 nm was measured per well.

### Coomassie gels

Plasma was diluted 1:50 in RIPA buffer (50 mM Tris pH 8.0, 150 mM sodium chloride, 1% NP-40, 0.5% sodium deoxycholate, 0.1% sodium dodecyl sulfate) with 5X Sample Buffer (250 mM Tris pH 6.8, 50% glycerol, 5% β-mercaptoethanol, 0.025% bromophenol blue, 5% SDS). Samples were separated on homemade 10% polyacrylamide gels then stained with Coomassie Blue for 30 minutes at room temperature with rocking. Gels were destained by incubating in water overnight. Gels were then imaged on an ImageQuant LAS 4000 luminescent image analyzer (GEHealthcare).

### Immunoblotting

To prepare protein lysates from plasma, plasma was harvested as described above and diluted 1:50 in RIPA buffer (50 mM Tris pH 8.0, 150 mM sodium chloride, 1% NP-40, 0.5% sodium deoxycholate, 0.1% sodium dodecyl sulfate) then combined with 5X sample buffer (250 mM Tris pH 6.8, 50% glycerol, 0.025% bromophenol blue, 5% sodium dodecyl sulfate, 5% beta-mercaptoethanol). Samples were separated on homemade 10% polyacrylamide gels and transferred to Immobilon-FL membranes (Millipore) via wet transfer. Membranes were blocked in 5% milk in TBST (50 mM Tris pH 8.0, 150 mM NaCl, 0.1% Tween-20) for 1 hour at room temperature. Membranes were incubated in primary antibody diluted in blocking solution at 4°C with rocking overnight and washed with TBST for 5 minutes five times. Membranes were incubated in HRP-conjugated secondary antibody diluted in blocking solution at room temperature with rocking for one hour and washed with TBST for five minutes five times. Membranes were incubated in ECL Prime Western Blotting Detection Reagent (GE Healthcare) for five minutes and imaged on an ImageQuant LAS 4000 imager (GE Healthcare). The primary antibody used was ALB (1:5,000, Abclonal #A24161). The secondary antibody was Rabbit (1:50,000, Abcam #ab205718).

### Global untargeted metabolomics

Untargeted metabolomics was performed by Metabolon, Inc. as previously described^29, 30^.

### Suppressive signal assay *ex vivo*

Master stocks of the following molecules were made fresh prior to each experiment: retinol (Cat.# AAJ62079MD) at 20 mM in 100% EtOH, L-Tryptophan (Cat.# T8941-25G) at 200 mM in DMSO, N-Formylanthranilic acid (Cat.# S636886-250MG) at 50 mM in DMSO. After harvesting hepatectomy plasma, it was split into separate 250 μL aliquots and treated with one of the following: retinol at 50 μM, L-Tryptophan at 1mM, N-Formylanthranilic acid at 100 μM, or the corresponding vehicle controls. The plasma and small molecule mixtures were incubated at 37°C with rotating for 75 minutes. After the incubation, all plasma was resuspended in 1mL EDTA K3E coated tubes from Sarstedt, then spun in Amicon® Ultra Centrifugal Filters with a 10 kDa MWCO to remove any unbound small molecules. Lastly, samples were again resuspended in 1mL EDTA K3E coated tubes from Sarstedt, volume brought up to 250 μL, then used for downstream assays.

### Displacement test *ex vivo*

Master stocks of the following molecules were made fresh prior to each experiment: palmitic acid (Cat.# P1145-5G) at 200 mM in 100% EtOH, oleic acid (Cat.# 031997.06) at 200 mM in 100% EtOH, linoleic acid (Cat.# L1012-1G) at 200 mM in 100% EtOH. After harvesting sham plasma, it was split into separate 250 μL aliquots and treated with one of the following: palmitic acid at 750 μM, oleic acid at 750 μM, linoleic acid at 750 μM, or the corresponding vehicle controls. The plasma and small molecule mixtures were incubated at 37°C with rotating for 75 minutes. After the incubation, all plasma was resuspended in 1mL EDTA K3E coated tubes from Sarstedt, then spun in Amicon® Ultra Centrifugal Filters with a 10 kDa MWCO to remove any unbound small molecules. Lastly, samples were again resuspended in 1mL EDTA K3E coated tubes from Sarstedt, volume brought up to 250 μL, then used for downstream assays.

### sgRNA design and cloning

sgRNA sequences were designed using the Broad Institute Genetic Perturbation Platform (GPP) CRISPick tool. Overlapping oligonucleotides containing CACC and AAAC 5’ overhangs were synthesized, phosphorylated with T4 polynucleotide kinase, annealed, and ligated into pLentiCRISPR vector linearized by Esp3I digestion and dephosphorylation with alkaline phosphatase. Ligations were transformed into Stbl2 cells and plated at 30°C to prevent recombination at LTRs. Plasmid was purified using ZymoPure kits with EndoZero columns to ensure endotoxin-free preparation.

### AAV preparation

AAV was produced as previously described^31^. Briefly, HEK293T/C17 cells were transfected with pAAV2/8, pHelper, and transgene plasmids using polyethylenimine (PEI). Viral supernatant was harvested at 72 h and again at 120 h post-transfection, and cells were lysed by scraping and resuspended in Salt-Active Nuclease (SAN) digestion buffer. Viral particles from the supernatant were concentrated by PEG precipitation, combined with the cell lysate, and purified by iodixanol density gradient ultracentrifugation (350,000G, 2.5 hours, 18°C). The 40% iodixanol fraction was collected and buffer-exchanged into DPBS using Amicon Ultra-15 centrifugal filter devices. Viral titers were determined using the Takara AAVpro Titration Kit.

### Administration of AAV to LSL-Cas9 mice

Mice were anesthetized with 2% isoflurane-oxygen delivered via nosecone. Mice were retroorbitally bled to acquire starting point of albumin concentration in the plasma. To infect the entire liver, mice were injected intraperitoneally with 4×10^11^ GC of either AAV-Cre-sgAlbumin or AAV-Cre-sgTigre.

### Organ-body weight measurement

Mice were weighed prior to anesthesia. Mice were bled as described above for plasma harvest. After harvesting plasma, each organ was removed from the mouse and rinsed in PBS. Each organ was individually weighed and calculated as percent of body weight.

## ACKNOWLEDGMENTS

We thank Bruce Enzmann for technical assistance and Jeffrey Kuhn of the Spatial Imaging Facility for microscopy assistance. We thank Stephen Bell and Matthew Vander Heiden for discussions. K.E.L. was supported by the NASEM Ford Foundation Predoctoral Fellowship, David H. Koch Graduate Fellowship, and MIT School of Science Graduate Fellowship. This work was supported by the National Cancer Institute Koch Institute Support Grant (P30CA014051).

## AUTHOR CONTRIBUTIONS

K.E.L. and K.A.K. conceived the project. K.E.L. performed all experiments and analyzed the data. K.E.L. and K.A.K. wrote the manuscript.

---

**Extended Data Figure 1.**
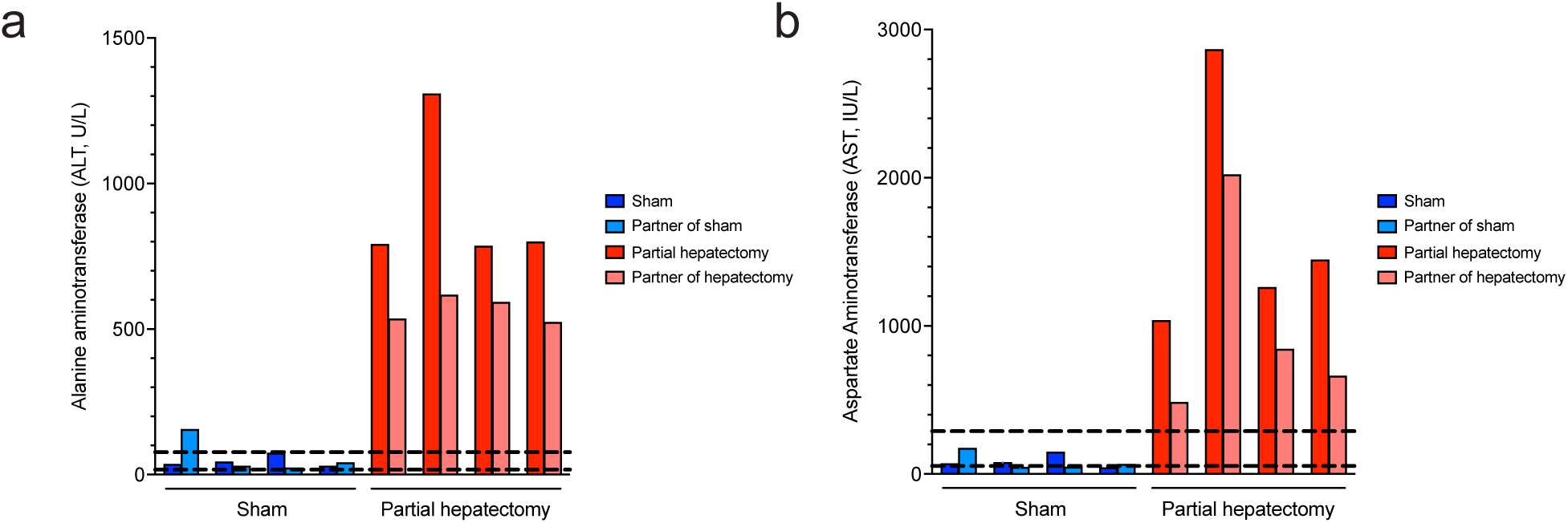
a) Measurements of ALT in serum of parabiotic mice 24 hours after partial hepatectomy or sham surgery. b) Measurements of AST in serum of parabiotic mice 24 hours after partial hepatectomy or sham surgery.

**Extended Data Figure 2:**
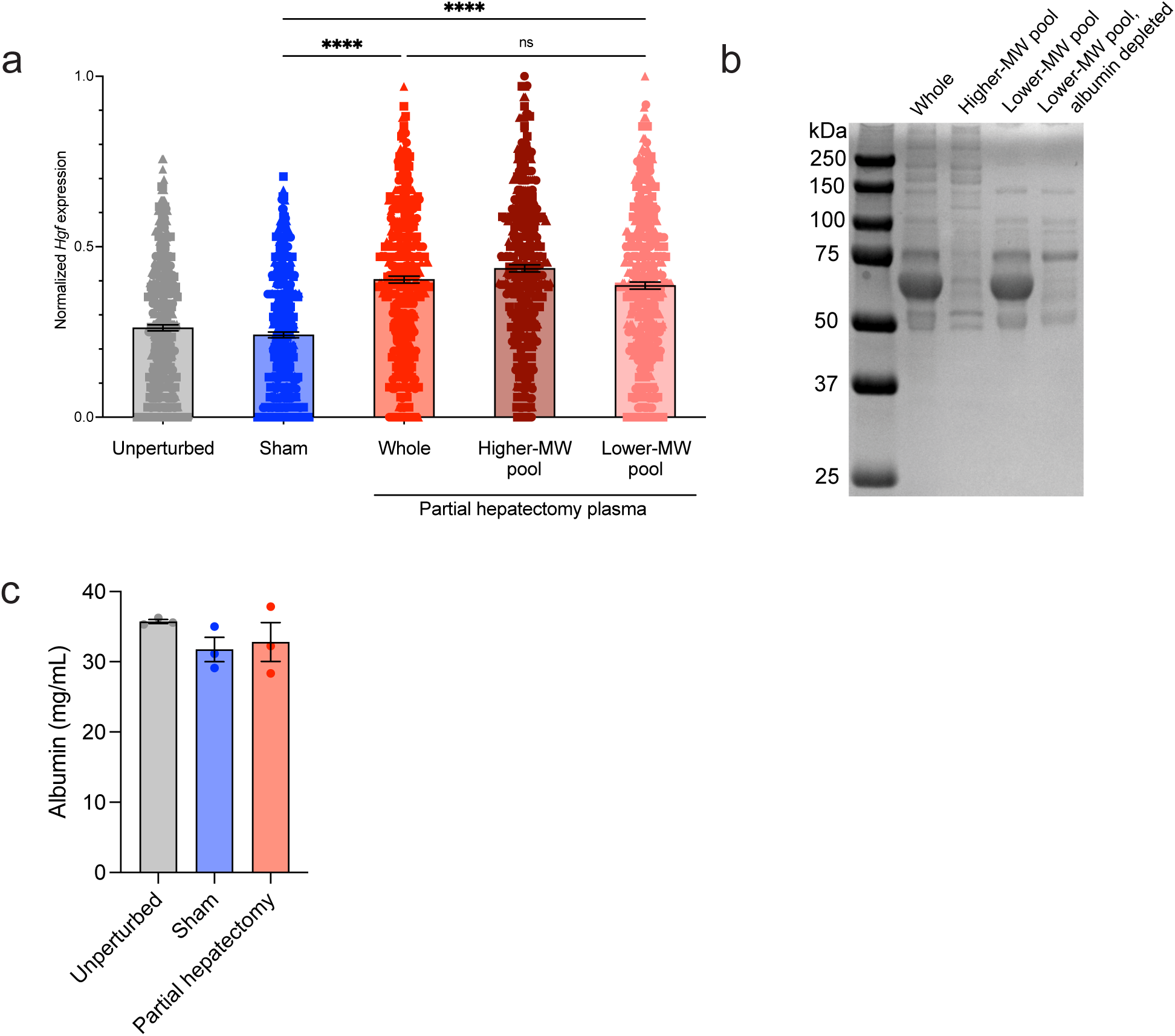
a) Normalized *Hgf* expression in stellate cells as determined by RNA FISH. n = 3 biological replicates with 150 cells analyzed per condition per replicate. Data are mean ± s.e.m. Sham versus whole p < 0.0001 and sham versus lower-MW pool p < 0.0001 by one-way ANOVA with Tukey’s multiple comparisons test. b) Coomassie-stained SDS-PAGE gel of equal input of whole unperturbed plasma, higher-, and lower-molecular weight (MW) pools and lower-molecular weight pool following albumin depletion. c) Albumin concentration in plasma harvested 12 hours after no perturbation, sham surgery, or partial hepatectomy as measured by ELISA. n = 3 biological replicates per condition. Data are mean ± s.e.m.

**Extended Data Figure 3:**
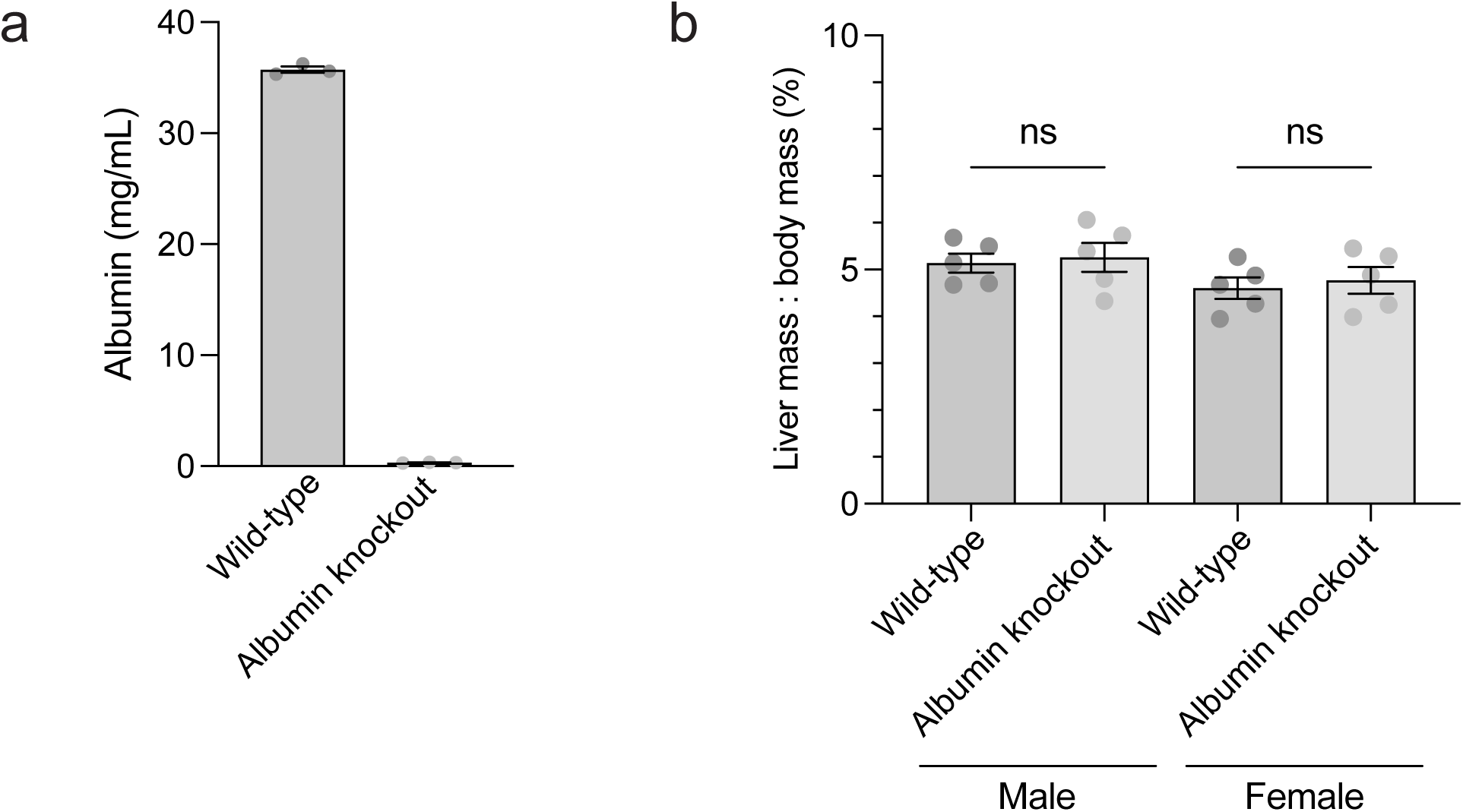
a) Albumin concentration in plasma harvested from unperturbed wild-type and albumin knockout mice as measured by ELISA. n = 3 biological replicates per genotype. Data are mean ± s.e.m. p < 0.0001 by two-tailed t test. b) Liver-to-body weight ratios for albumin knockout mice and wild-type littermates. Data are mean ± s.e.m.

**Extended Data Figure 4:**
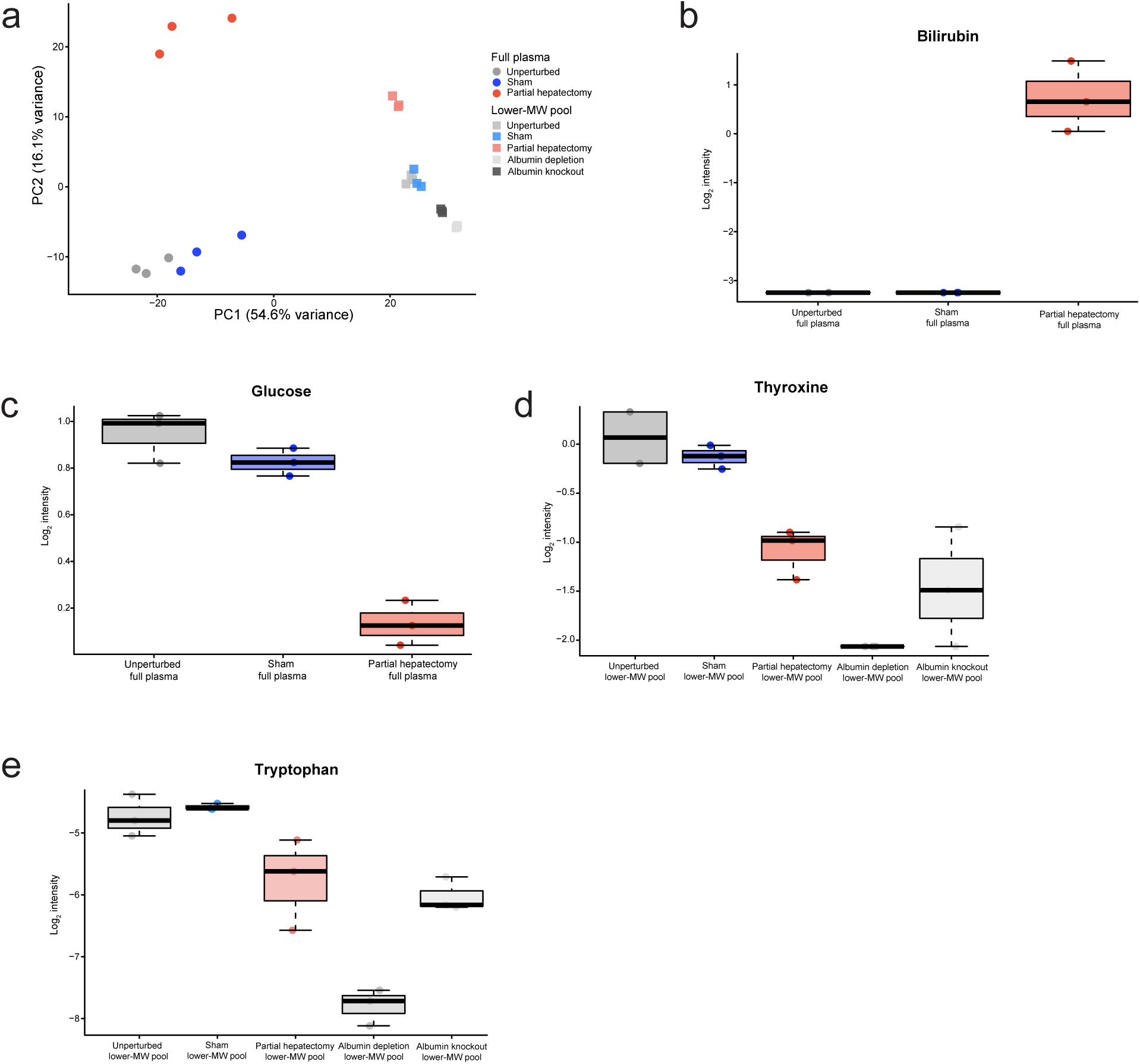
a) PCA plot of all samples submitted for untargeted metabolomics. b) Boxplot of relative abundance of bilirubin in full unperturbed, sham, or hepatectomy plasma. Boxplots show the median (center line), interquartile range (box), and 1.5× IQR whiskers. Points represent individual samples. c) Boxplot of relative abundance of glucose in full unperturbed, sham, or hepatectomy plasma. Boxplots show the median (center line), interquartile range (box), and 1.5× IQR whiskers. Points represent individual samples. d) Boxplot of relative abundance of thyroxine in unperturbed, sham, hepatectomy, albumin depleted, and albumin knockout lower-molecular weight pools. Boxplots show the median (center line), interquartile range (box), and 1.5× IQR whiskers. Points represent individual samples. e) Box plot of relative abundance of tryptophan in unperturbed, sham, hepatectomy, albumin depleted, and albumin knockout lower-molecular weight pools. Boxplots show the median (center line), interquartile range (box), and 1.5× IQR whiskers. Points represent individual samples.

**Extended Data Figure 5:**
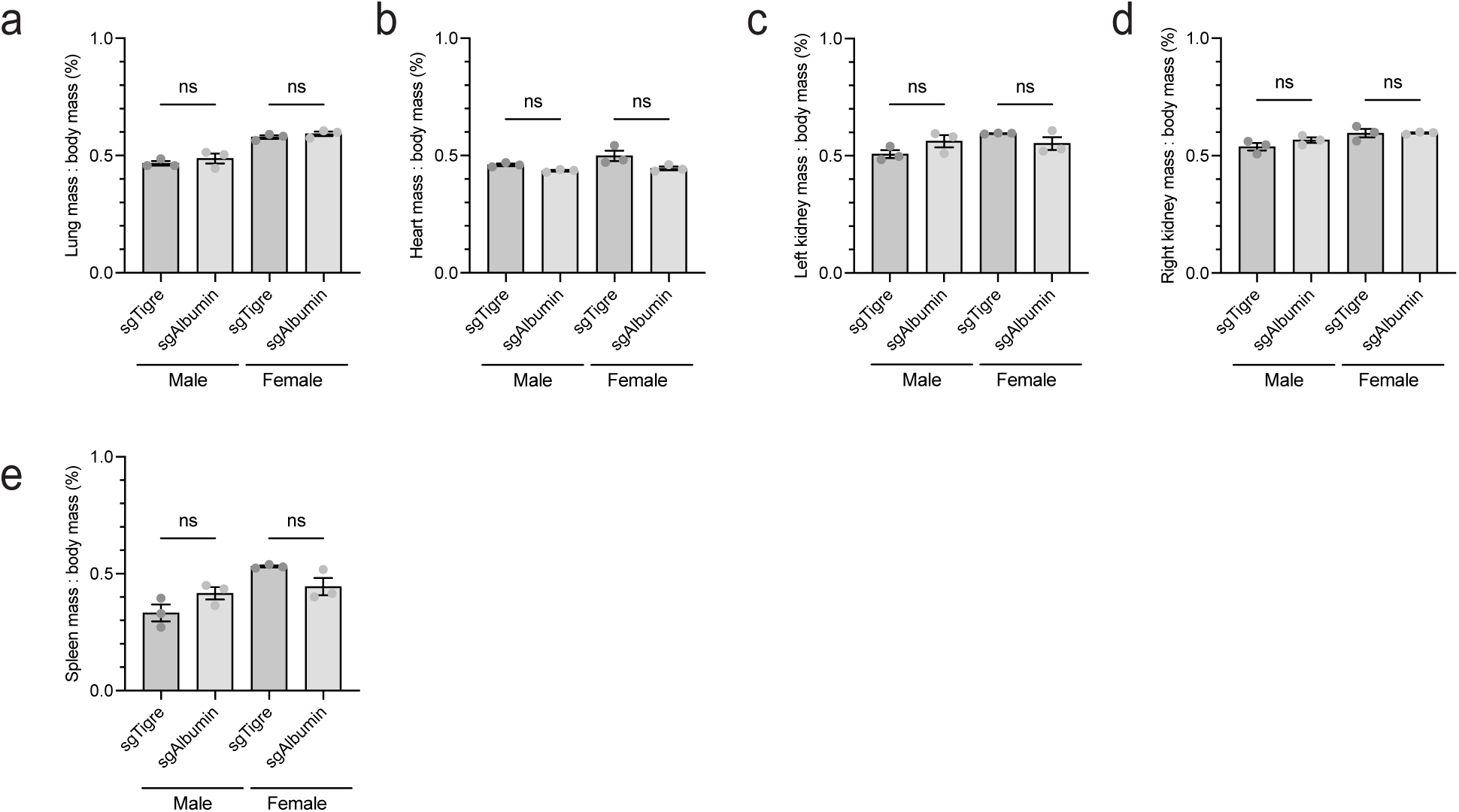
a) Lung-to-body weight ratio 56 days after injection with AAV-Cre-sgTigre or AAV-Cre-sgAlbumin. Data are mean ± s.e.m. b) Heart-to-body weight ratio 56 days after injection with AAV-Cre-sgTigre or AAV-Cre-sgAlbumin. Data are mean ± s.e.m. c) Left kidney-to-body weight ratio 56 days after injection with AAV-Cre-sgTigre or AAV-Cre-sgAlbumin. Data are mean ± s.e.m. d) Right kidney-to-body weight ratio 56 days after injection with AAV-Cre-sgTigre or AAV-Cre-sgAlbumin. Data are mean ± s.e.m. e) Spleen-to-body weight ratio 56 days after injection with AAV-Cre-sgTigre or AAV-Cre-sgAlbumin. Data are mean ± s.e.m.

